# PEDOT:PSS conducting eutectogel for enhanced electrical recording and stimulation in implantable neural interfaces

**DOI:** 10.1101/2025.09.14.674390

**Authors:** Martin Kasavetov, Rubén Ruiz-Mateos Serrano, Antonio Dominguez-Alfaro, Chaeyeon Lee, Matias L. Picchio, David Mecerreyes, George G. Malliaras, Viviana Rincón Montes, Amparo Güemes González

## Abstract

Conductive polymers such as PEDOT:PSS are widely used in bioelectronic interfaces due to their mixed ionic-electronic conductivity and biocompatibility. However, their mechanical fragility and limited processability constrain their performance in implantable devices. Deep eutectic solvents (DES), when combined with PEDOT:PSS, form eutectogels that enable thick, soft coatings. Here, we present a PEDOT:PSS-based eutectogel incorporating choline chloride: lactic acid and GOPS, integrated into flexible thin-film electrode arrays for sciatic nerve interfacing. These implants feature an array of electrodes and a pre-formed spiral geometry to conformally wrap small-diameter nerves. Devices were fabricated using standard photolithography and reactive ion etching techniques, allowing side-by-side comparison of PEDOT:PSS/DES with conventional PEDOT:PSS electrodes. PEDOT:PSS/DES enabled single-layer films up to 800 nm thick, significantly greater than PEDOT:PSS, and yielding over two-fold improvements in impedance and charge injection capacity *in vitro*. Acute *in vivo* electrophysiology in rats confirmed enhanced neural recording and stimulation capabilities, with lower impedance, higher capacitance, and reduced motor activation thresholds. While PEDOT:PSS/DES more reliably elicited motor responses at lower stimulation currents, electromyogram signal amplitudes from the tibialis anterioris at matched stimulation levels were comparable between materials. These results suggest that while superior electrochemical properties improve neural interface performance, local electrode-tissue interactions remain critical. Overall, this work establishes DES-modified PEDOT:PSS as a promising electrode material for soft neural interfaces and highlights its potential for advancing implantable bioelectronics.

## Introduction

Implantable bioelectronic devices have shown significant promise in treating a variety of pathological conditions, including arrhythmias, deafness, drug-resistant epilepsy, and Parkinson’s disease, among others^1–5^. In particular, thin-film microelectrode arrays have enabled detailed study and monitoring of both the central and peripheral nervous systems^6–13^. The principal advantage of these devices lies in their ultra-thin profiles, often just a few microns thick, which allows for excellent conformability, minimal foreign body reaction, and close contact between the electrodes and neural tissue^6,14^. Poly(3,4-ethylenedioxythiophene) doped with polystyrene sulfonate (PEDOT:PSS) is a material that has gained popularity in the last decades as an electrode coating for thin-film electrode arrays due to its exceptional mixed ionic-electronic conductivity and volumetric capacitance^15^. These properties enable the fabrication of small, high-performance electrodes with low impedance and mechanical flexibility, while maintaining biocompatibility and stability during electrical stimulation^16–18^.

Despite its improved characteristics, its application in implantable neurotechnology remains constrained by several critical limitations. Thin-film PEDOT:PSS coatings, especially when electrodeposited or spin-coated, are mechanically fragile and prone to delamination^19–21^, particularly under mechanical stress or during chronic implantation. This is a major concern for long-term stability *in vivo*, where micromotion between the implant and tissue is inevitable. Recent studies have demonstrated that electrodes based on conventional PEDOT:PSS formulations, including additives such as ethylene glycol (EG) and dodecylbenzene sulfonic acid (DBSA), exhibit superior stability compared to both Au and PEDOT:PSS-coated Au electrodes under accelerated aging conditions involving electrical stimulation, oxidative stress, and mechanical agitation^22^. These findings highlight the potential of PEDOT:PSS as a standalone material for chronic neuromodulation applications. PEDOT:PSS also offers significantly higher charge injection capacity (CIC) than conventional metal coatings^20,23,24^, making it attractive for electrical stimulation. However, further enhancement is needed to meet the demands of high-resolution stimulation and the ongoing miniaturisation of neural probes. Despite these advantages, PEDOT:PSS is typically processed as a thin film. Single spin-coated layers typically achieve thicknesses of 80–300 nm^21,25,26^, and while thicker films can be fabricated by stacking multiple layers, such multilayer structures often suffer from poor mechanical integrity and increased risk of delamination^21,26^. Its limited processability into thicker films therefore restricts the performance of electrodes. This is suboptimal for interfacing with soft, curved, and dynamic neural tissues, such as small peripheral nerves and the enteric nervous system, where conformability and mechanical integration are essential for stable, long-term performance ^9,11,27^

To overcome these limitations, deep eutectic solvents (DES) have emerged as a promising additive to impart printability of three-dimensional (3D) coatings and enhance properties of PEDOT:PSS^28^. DES are a novel class of ionic additives, similar to ionic liquids, formed by mixing two or more components, typically a hydrogen bond donor (HBD) and a hydrogen bond acceptor (HBA), that interact through hydrogen bonding to produce an eutectic mixture with an abnormal depression in its melting point^29–32^. These mixtures are liquid in a broad temperature range, can be easily synthesised through green, low-cost processes, and exhibit low toxicity^33^, making them attractive for biomedical applications. When mixed with PEDOT:PSS, DES act as gelators, forming supramolecular networks denoted as conducting eutectogel. These gels possess shear-thinning properties suitable for extrusion-based 3D printing^34^, enabling the fabrication of soft, conformable electrode architectures. In addition to enhancing processability, DES significantly improve both ionic and electronic conductivity compared to pristine PEDOT:PSS and plasticise the polymer at low concentrations, increasing its self-standability^35–39^. Altogether, DES incorporation enables the fabrication of mechanically robust, self-supporting 3D electrode architectures, representing a significant step forward in addressing the structural fragility and integration challenges of thick conventional PEDOT:PSS coatings^40–47^. Nonetheless, the rapid gelation of such materials^34^ could pose challenges for their applicability in thin film technology. To the best of our knowledge, conducting eutectogels based on PEDOT:PSS and DES have never been reported in the literature for implantable bioelectronics.

In this work, we build upon a previously reported formulation of PEDOT:PSS combined with the DES formed by choline chloride and lactic acid (ChCl:LAC), by introducing an optimised concentration of (3-glycidyloxypropyl)trimethoxysilane (GOPS) as crosslinker. The addition of GOPS is critical to ensure the mechanical and chemical robustness of the material during processing, thereby enabling the microfabrication of thin-film electrode arrays. We extensively evaluated the resulting PEDOT:PSS/ChCl:LAC/GOPS electrodes, hereafter referred to as PEDOT:PSS/DES, and compared against PEDOT:PSS electrodes using electrochemical and electrical characterizations in an *in vitro* setting to validate the performance of the materials for neural recording and stimulation Finally, we validated the functionality and overall improved capabilities of the electrodes in acute *in vivo* experiments by electrically recording and stimulating the sciatic nerve in rats. Our results demonstrate that the PEDOT:PSS/DES formulation yields thicker films, resulting in enhanced electrochemical properties than conventional PEDOT:PSS formulations under identical spin-coating conditions, supporting it as a high-performance material platform for implantable bioelectronic interfaces.

## Experimental

### DES synthesis

DES were synthesised using the thermal mixing method. ChCl and LAC were combined at a molar ratio of 1:2 and heated to 60 °C under constant stirring until a clear, homogeneous liquid was formed.

### Preparation of PEDOT:PSS/ChCl:LAC/GOPS eutectogels

PEDOT:PSS (Clevios™ PH1000) and the synthesised DES (ChCl:LAC) were mixed at room temperature, resulting in an estimated DES concentration of 1.3 % (w/w) in PH1000. The mixture was vortexed for approximately 20 seconds to ensure homogeneity. Next, GOPS was added to with a concentration of 0.65% (w/w). The solution was vortexed again for 20 seconds. Upon complete mixing, the composite exhibited a noticeable increase in viscosity, forming a gel-like consistency. The resulting eutectogel was loaded into a syringe, filtered using a polyvinylidene difluoride (PVDF) filter with a pore size of 0.4 µm, and deposited onto wafers for further processing.

### Preparation of conventional PEDOT:PSS

The conventional PEDOT:PSS formulation was prepared similar to previously published protocols^48^. Briefly, PEDOT:PSS (Clevios™ PH1000) was mixed with 5% (v/v) of EG and 0.25% (w/w) of DBSA and subjected to ultrasonication for 10 minutes. Next, 1% (v/v) of GOPS was added to the mixture, followed by an additional 1 minute of ultrasonication. Prior to deposition of the wafer, the solution was filtered using a PVDF filter with a pore size of 0.4 µm.

### Device fabrication

The microfabrication was carried out at the Helmholtz Nano Facility at the *Forschungszentrum* Jülich^49^. The devices were fabricated using conventional microfabrication techniques similar to previously reported protocols^50^ and feature a layer of metal and a conductive polymer film (PEDOT:PSS/DES or PEDOT:PSS) embedded between two insulating layers of parylene-C (PaC).

As a first step, 2 µm of PaC were deposited on a 4-inch silicon host wafer *via* chemical vapor deposition (CVD) using the PDS 2010 Labcoater 2 tool (Specialty Coating Systems Inc., USA), followed by deposition of the metal layer. The latter was patterned using a bilayer lift-off mask that comprised a LOR3B (Micro Resist Technology, Germany) bottom layer and an AZ nLOF2020 (MicroChemicals GmbH, Germany) top layer as image resist. Prior to spin-coating, the PaC substrate was dehydrated for 2 minutes at 120°C using a direct contact hotplate. The LOR3B layer was spin coated at 3000 rpm for 30 seconds with a ramp of 500 rpm/sec and soft-baked at 150°C for 5 minutes on a direct contact hotplate. After cooling down, the AZ nLOF2020 layer was spin-coated using the same parameters, followed by a final soft bake at 120°C for 1 minute. The photoresist was exposed using a maskless aligner (MLA-150, Heidelberg Instruments, Germany) with a dose of 260 mJ/cm^2^ and a defocus distance (defoc) of 0 at a wavelength of 375 nm, followed by a post-exposure bake (PEB) at 110°C for 2 minutes. The lift-off mask was finally developed in AZ 326 MIF (MicroChemicals GmbH, Germany) for 35 seconds and rinsed in a deionized water cascade.

The metal layer, comprising 20 nm of titanium (Ti) and 100 nm of gold Au), was deposited *via* electron-beam assisted evaporation using the Univex 400 physical vapor deposition tool (Leybold GbmH, Germany) with a deposition rate of 0.1 nm/s for Ti and 0.5 nm/s for Au. For the lift-off step, the wafers were submerged in acetone for 2-3 hours until the metal residues were lifted and rinsed in isopropanol. Finally, the solvent-resistant LOR3B layer was stripped in AZ 326 MIF for 5 minutes, followed by a rinse in a water cascade.

The conductive polymer film was deposited after patterning the metal layer *via* spin-coating. Prior to spin-coating, the wafers were dehydrated for 2 minutes at 120°C and subjected to oxygen plasma activation at 0.8 mbar O^2^ with a power of 80 W for 3 minutes (Gigabatch 310M, PVA TePla AG, Germany). For each wafer, 10 ml of PEDOT:PSS or PEDOT:PSS/DES was prepared as described previously and spin-coated at 500 rpm with a ramp of 5000 rpm/s for 60 seconds. The resulting films were hard-baked at 120°C for 1 hour on a direct contact hotplate. The PEDOT:PSS film was additionally soaked in deionized water overnight (~12 hours) to induce swelling and remove the volatile components.

Both the PEDOT:PSS and PEDOT:PSS/DES films were patterned *via* reactive ion etching (RIE) using a CF4:O2 (5:50 sccm) gas mix and RF power of 150 W (Oxford Plasmalab 100-ICP 180, Oxford Instruments, United Kingdom) using AZ5214E as a positive photoresist etch mask. The latter was prepared by spin-coating at 3000 rpm (PEDOT:PSS) or 2500 rpm (PEDOT:PSS/DES) for 30 seconds after initial 5 seconds at 500 rpm with an acceleration of 500 rpm/s, followed by soft-baking at 110°C for 1 min. The positive photoresist was exposed with a dose of 100 mJ/cm^2^ and a defoc of 0 using the MLA-150 and developed in AZ 326 MIF for 70 seconds. In the case of PEDOT:PSS, the photoresist was spin-coated directly on top of the conductive polymer film, while PEDOT:PSS/DES films were first coated with a 100 nm-thick layer of PaC to promote the adhesion of the etch mask. This interlayer was etched through during the PEDOT:PSS/DES etching step, facilitating the structuring of the conductive polymer film. Lastly, the etch mask was stripped in acetone using gentle ultrasonication and rinsed in isopropanol.

To passivate the devices, an additional 2 µm of PaC were deposited as described before^48^. Silane A127 adhesion promoter was applied in the chamber using the swab method to facilitate the adhesion of the passivation layer. Afterwards, the outline of the devices, as well as the openings of the electrodes and contact pads were patterned using RIE in two steps. To this end, a 15 µm-thick positive photoresist layer of AZ12XT (MicroChemicals GmbH, Germany) was spin-coated at 1000 rpm for 180 seconds with an acceleration of 200 rpm/s and soft-baked at 100°C for 4 minutes. The resist was subsequently exposed with a dose of 350 mJ/cm^2^ and a defoc of 2 with maskless lithography (MLA-150), subjected to a PEB (1 min at 90°C) and developed in AZ 326 MIF for 2 minutes. The outline of the probes, including the openings of the metal contact pads, was etched in the first step using a CF4:O2 (4:36 sccm) gas mix, RF power of 50 W, and ICP power of 500 W. The etch mask was stripped using two baths of AZ 100 Remover, followed by one bath of acetone and one bath of isopropanol, each comprising two to three minutes. Gentle ultrasonication was applied in the first bath with AZ 100 Remover to aid the dissolution of the photoresist. The wafers were finally dried using a compressed nitrogen gun. After stripping the resist, a new layer of AZ12XT was spin-coated, and the aforementioned process was repeated to etch the electrode openings.

This process flow was used to fabricate probes for both *in vitro* and *in vivo* settings. After microfabrication, the *in vitro* probes were released from the host substrate using drops of deionised water. They were then flip-chip bonded manually on custom-made printed-circuit boards (PCBs) using a low-temperature solder paste alloy (SMDLTB35T4, Chip Quik, Sn60/Bi40) at 180°C and cooled down to room temperature. Conversely, the *in vivo* probes were bonded directly on the host wafer. A customised flat flexible cable (FFC)^51^ was bonded using anisotropic conductive film (ACF, TESA, Germany) and a FINEPLACER pico2 (Finetech GmbH &Co. KG, Germany), as previously described^48^. After bonding, the flexible probes were released from the wafer using drops of deionised water.

### Electrochemical and electrical characterization

All electrochemical measurements were carried out in a bath of saline solution (1xPBS). Electrochemical impedance spectroscopy (EIS) was performed with the VSP-300 (BioLogic Science Instruments, France) potentiostat using a 10-mV sinusoidal signal and a frequency sweep from 1 Hz to 1 MHz. A 3-electrode setup was utilised comprising a Ag/AgCl reference electrode, a Pt wire as a counter electrode, and the device electrodes serving as individual working electrodes. To further evaluate the performance of the electrodes, the impedance magnitude (|*Z*|) was extracted at 1 kHz as well as the specific capacitance (*C*_*s*_) from the EIS data. The specific capacitance was estimated according to Eq. 1:

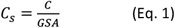

where *GSA* is the geometric surface area of the electrodes and *C* is the equivalent capacitance, extracted by performing a *Z* fit at the low-frequency regime (1 Hz – 100 Hz) using a simple Randles circuit^52^. In addition, we estimated the thermal noise level (υ_*n*_) in the relevant frequency bands according to Eq. 2^53^:

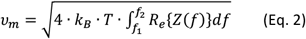

where *k*_*B*_ is the Boltzmann constant, *f*_*1*_ and *f*_*2*_ are the lower and upper limits of the frequency band of interest (*f*_*1*_ = 300 Hz and *f*_*2*_ = 3 kHz for the higher frequency band, and *f*_*1*_ = 1 Hz and *f*_*2*_ = 300 Hz for the lower frequency band), *T* = 300 K is the temperature, and *R*_*e*_{*Z(f)*} is the real component of the impedance.

To estimate the cathodic charge storage capacity (*CSC*_*c*_), we performed cyclic voltammetry between the potential limits of −0.9 V and 0.6 V, corresponding to the water window for conductive polymers^54^. The measurements were carried out on a multichannel potentiostat (CH Instruments Inc., USA) in a 3-electrode setup as described above with a scan rate of 100 mV/s and a current sensitivity level of 100 nA. At least 5 cycles were carried out for each electrode and only the last cycle was taken for further analysis. The cathodic *CSC*_*C*_ was estimated by integrating the cathodic current *I*_*E*_ over the potential limits (*E*_*a*_=0.6 V and *E*_*c*_=-0.9 V) according to Eq. 3, where υ is the scan rate:

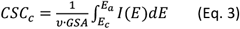

To estimate the *CIC*, we utilized the current-controlled stimulator of the ME2100 system (Multi Channel Systems MCS GmbH, Germany) to deliver current pulses of defined amplitude and duration to the electrodes, while recording the resulting voltage transients with a 2208 Pico-Scope oscilloscope (Pico Technology, UK). Measurements were performed in saline solution using a two-electrode setup, with an Ag/AgCl pellet serving as both reference and counter electrode, and the oscilloscope connected in parallel to a working electrode. We utilized square biphasic charge-balanced pulses with a leading cathodic phase, followed by a symmetrical anodic phase with an interphase gap of 20 µs. For the 100 µm size electrodes, which were implemented in the *in vivo* design, we tested four different phase durations (0.1 ms, 0.5 ms, 1 ms, and 5 ms) in order to gain insights into the time-dependent properties of the charge injection mechanism. To account for size-dependent scaling effects, we additionally measured the CIC for smaller electrodes (25 and 50 µm) at a fixed phase duration of 0.5 ms. In each measurement, the phase duration (*T*_*ph*_) and pulse shape were kept constant, and the current amplitude was gradually increased until the polarization limit of the electrodes was reached. As explained earlier, we assumed polarization limits of −0.9 V and 0.6 V for the cathodic and anodic regimes, respectively. The maximum voltage excursion during each phase of the pulse was extracted by subtracting the access voltage (*V*_*A*_) and compared with the anodic and cathodic potential limits of the electrodes. In all measurements, the cathodic polarization limit was reached first. After extracting the maximum current at the polarization limit (*I*_*inj,max*_), the *CIC* of each electrode was calculated according to Eq. 4:

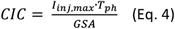

The sheet resistance of the films was measured in dry conditions at room temperature with the Van der Pauw (VDP) method using a Keithley 4200 with probe station MPI-TS200 (Tektronix Inc., USA) and a 4-probe setup using tungsten probes (Picoprobes ST-20-2). To facilitate the measurements, we fabricated specialised test structures to aid the contacting of the conductive polymer film. The test structures comprised metal pads (Ti/Au) contacting the edges of a 500 µm wide square of conductive polymer film. The structures were fabricated on a PaC-coated wafer with the process described earlier, and the conductive polymer squares were structured *via* dry etching. The conductivity of the films was estimated from the sheet resistance and the film thickness using Eq. 5:

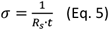

where *R*_*s*_ is the sheet resistance and *t* is the film thickness. The thickness measurements were performed using a Dektak XT profilometer (Bruker, USA) with a vertical resolution of 2 nm. The thickness in the finalized devices was estimated by focused ion-beam (FIB) cuts and scanning electron microscopy (SEM) of the electrode cross section using an FEI TFS Helios 600i (G3) FIB/SEM dual-beam instrument.

### Statistical analysis for *in vitro* comparisons

Statistical analysis was performed using self-written scripts in Python utilizing the *statistics* package. Unless stated otherwise, the data is depicted as mean ± standard deviation. Normality of the data was tested using a Shapiro-Wilk test. If normality was rejected, a Mann-Whittney U test was employed to compare the data.

### Surgery & *in vivo* electrophysiology measurements

All experimental procedures were performed in accordance with the UK Animals (Scientific Procedures) Act 1986 and were approved by the animal welfare ethical review body at the University of Cambridge. These procedures were performed under a project license (PP5478947) by A. Güemes (personal licence I10076024), issued by the UK Home Office. Non-recovery surgeries were performed in four female Sprague-Dawley rats (purchased at 200 - 250 g) (Charles River Laboratories, Kent, UK) with bilateral sciatic nerve implantation, resulting in a total of eight nerves (*N*=8). Animals were anaesthetized with urethane (1.2–1.5 g/kg, intraperitoneally), and anaesthesia depth was monitored throughout. Body temperature was maintained using a heating pad, and the exposed nerve was kept moist with sterile saline throughout the experiment.

A mid-thigh incision was made to expose the sciatic nerve. Two types of cuff electrode coatings, PEDOT:PSS/DES and PEDOT:PSS, were implanted side by side on each nerve prior to its bifurcation, with their positions alternated across animals to control for proximal-distal effects. Care was taken to ensure consistent placement of corresponding electrodes from both cuffs around the nerve to enable a fair comparison.

Neural and EMG signals were acquired using the Intan RHS system (RHS2000/Stim, Intan Technologies) at a sampling rate of 30 kHz. The PEDOT/DES and PEDOT:PSS cuffs were connected to two separate ports to allow simultaneous recording, and the EMG needle was connected to an additional port. Two ground electrodes, connected to the respective GND/reference input of the headstage, were placed subcutaneously in two small pockets, one for the neural electrodes and another for the EMG electrode. For EMG recordings, the skin over the tibialis anterior muscle was incised to expose the muscle belly, and a 30G needle electrode was inserted directly into the muscle.

Each nerve underwent an electrophysiology protocol lasting approximately 1 hour. Spontaneous and pain-evoked activity was recorded first, beginning with a 5-minute baseline followed by repeated mechanical stimulation of the tibialis anterior muscle using a needle press to evoke nociceptive responses. This was followed by electrical stimulation through the implanted cuffs in monopolar configuration. Stimulation was delivered using single pulses (100 µs pulse width) at increasing amplitudes: from 5 µA to 10 µA in 1 µA steps, and then at 15, 20, and 30 µA. Each stimulation trial consisted of a 10-second baseline followed by 10 pulses at 1 Hz (1-second inter-pulse interval, pre-set in Intan) stimulating with one cuff, followed by 10 pulses stimulating with the other cuff at 1Hz, alternating between cuffs to compare responses. Channels with impedance values exceeding 10 kΩ were excluded from further analysis.

### Analysis of electrophysiology signals

Impedance magnitude and phase values were extracted from the Intan file metadata corresponding to the baseline recording for each available circular channel (100 µm diameter). From these values, the equivalent series resistance (*R*) and capacitance (*C*) of each electrode were computed using standard impedance modelling techniques, assuming a series RC circuit approximation.

Offline data processing was performed in Python, where signals were bandpass filtered between 200-3000 Hz for high-frequency activity analysis. For the analysis of pain-evoked activity, a 60-second segment of baseline activity and a 60-second segment during mechanical stimulation were selected. The root mean square (RMS) of the signal was computed for each channel, and the values were averaged across channels. Signal-to-noise ratio (SNR) was calculated as the ratio of RMS during evoked activity to RMS during baseline. Spikes elicited by the stimulation were detected using a negative threshold of −12 µV. Waveforms were extracted using a 4-ms window centred around the negative peak and averaged to characterize the evoked response. The same procedure was applied to the capsaicin and nerve-press recordings.

For electrical stimulation analysis, stimulation artifacts were detected using an adaptive threshold, with a minimum inter-artifact interval of 1 second to match the stimulation frequency. A post-stimulus window from 11 ms to 200 ms after each artifact was extracted to capture evoked neural activity while avoiding contamination from the artifact itself, based on expected conduction velocities. RMS values were computed within this window and averaged across trials. Baseline RMS was also calculated using a window of the same duration immediately preceding the onset of stimulation. The increase in RMS relative to baseline was compared between the two electrode materials. Additionally, the stimulation threshold required to elicit a visible muscle twitch was noted during experiments and compared across materials.

Population-level analysis was performed using paired statistical comparisons within each animal. Assumptions were checked prior to applying parametric t-tests by testing whether the differences between paired values were normally distributed using the Shapiro-Wilk test. If normality was violated, the non-parametric Wilcoxon signed-rank test was used as an alternative. The number of available channels for each material varied between 4 and 9 in all trials, but was always kept consistent between the two recording ports within each trial to ensure fair pairwise comparison between material types. For the analysis of impedance and equivalent RC, values were averaged across available channels for each nerve, and paired statistical comparisons were made between the two materials. To visualize variability and distribution, boxplots were also generated for all individual channel impedance and RC equivalent values across all trials and animals. For the recording and stimulation-related metrics (RMS, SNR, muscle stimulation threshold), the difference between the two materials (Δ) was computed for comparison purposes, and paired statistical comparisons were also assessed.

## Results and discussion

### Material formulation, device design and fabrication

To explore the potential of PEDOT:PSS eutectogel enhanced with DES, we selected a composition using lactic acid (LAC) as the HBD and choline chloride (ChCl) as the HBA (Figure 1a, left). This specific DES (ChCl:LAC) was chosen based on prior studies demonstrating improved conductivity in thick pristine PEDOT:PSS films deposited *via* drop-casting^55^. We formulated a PEDOT:PSS/DES blend by mixing 1.3% (w/w) of DES with PH1000, followed by the addition of 0.65% (w/w) of GOPS, resulting in a PEDOT:DES:GOPS weight-ratio of 2:2:1. The addition of GOPS to the blend results in cross-linking of the film during the hard baking step, enhancing the stability of the material and thereby ensuring compatibility with standard microfabrication techniques.

**Figure 1.**
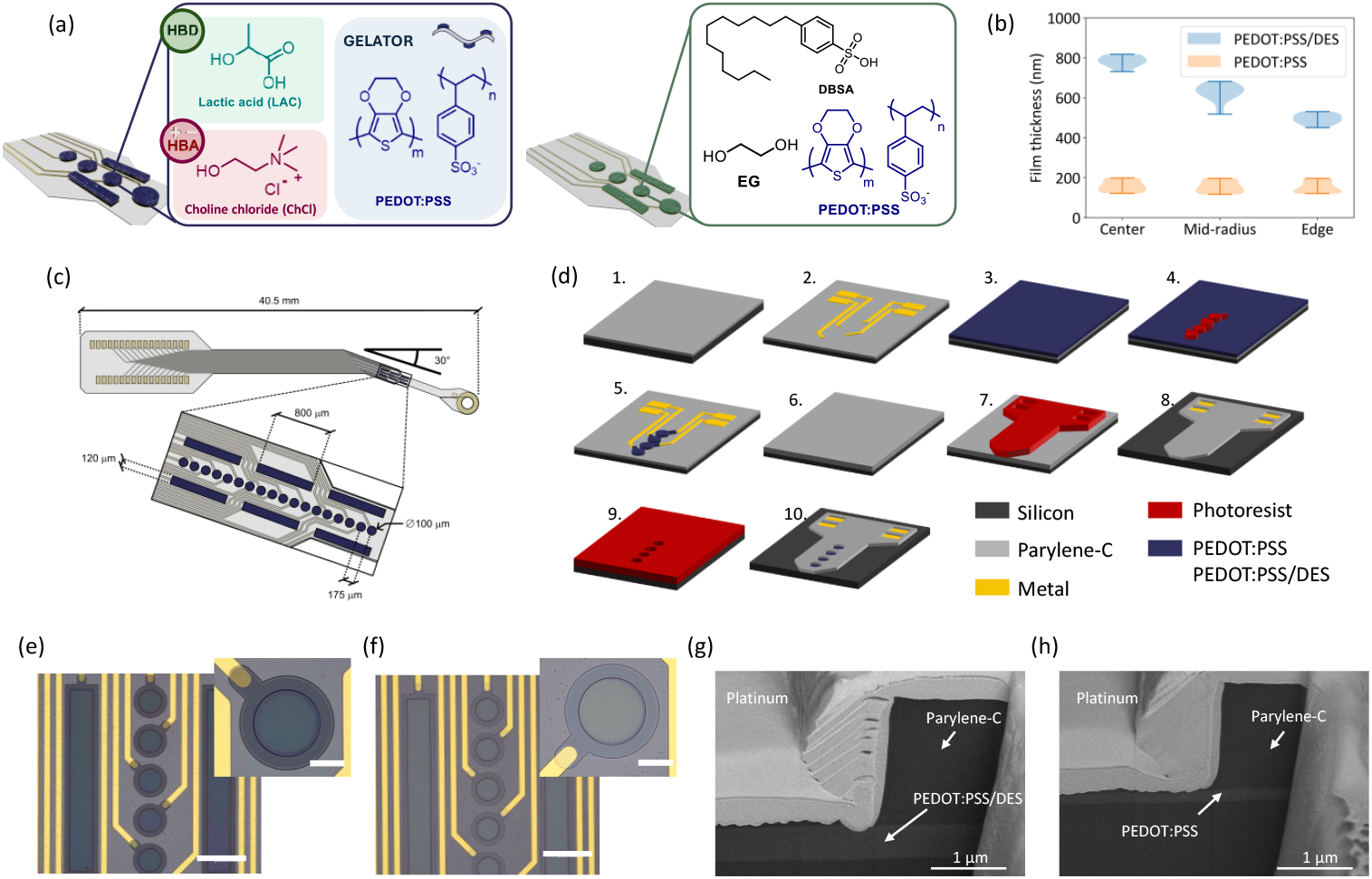
Material formulation, device design and fabrication. (a) Material composition of the PEDOT:PSS/DES blend proposed in this work *(left)*. The formulation combines PEDOT:PSS, DES and GOPS at a weight ratio of 2:2:1. The performance of the new blend is compared to a standard PEDOT:PSS formulation *(right)* featuring EG, GOPS, and DBSA. (*b*) Violin plots illustrating the comparison of the wafer distribution of the film thickness of spin-coated films with the PEDOT:PSS/DES blend (blue) and the conventional PEDOT:PSS blend (orange). PEDOT:PSS/DES films exhibit a reduction in thickness towards the edge of the wafer. (*c*) Schematic of the *in vivo* probe used in this work. The design is adapted for interfacing with the rat sciatic nerve. (*d*) Process flow describing the fabrication of the devices. The process comprises the following steps: (1) CVD deposition of a PaC substrate layer (2 µm) on a 4” Si carrier wafer, (2) E-beam evaporation of the metal layer (Ti/Au) and patterning *via* lift-off. The metal layer comprises the feedlines as well as the contact pads for the front-end connector. (3) Spin-coating of the conductive polymer film and patterning of the film *via* RIE (4 and 5) to form metal-free electrodes. Deposition of a PaC encapsulation layer (6) with a thickness of 2 µm. Patterning of the shape and electrode openings *via* RIE (7 and 8) followed by etching of the electrode openings (9 and 10). (*e, f*) Optical microscopy pictures of finalized *in vivo* probes, highlighting the sensing area of the devices. PEDOT:PSS/DES (*e*) devices feature an enhanced film thickness compared to conventional PEDOT:PSS devices (*f*). Insets show a close-up of the metal free polymeric electrodes. Scale bars correspond to 200 µm (main images) and 50 µm (insets). *(g, h*) SEM images of the cross section of 25 µm PEDOT:PSS/DES (*g*) and PEDOT:PSS (*h*) electrodes obtained *via* FIB cuts. The estimated film thickness at the edge of the electrodes is 550 nm for the PEDOT:PSS/DES and 171 nm for the PEDOT:PSS electrode.

Conventional PEDOT:PSS formulations often contain additives (Figure 1a, right) such as ethylene glycol (EG) and dodecylbenzene sulfonic acid (DBSA) to enhance conductivity *via* conformational changes in PEDOT chains, from coiled to linear structures, driven by reduced water evaporation and a screening effect between the PSS-rich polymer and its dopant^56,57^. In contrast, DES-based additives achieve similar or superior performance. During evaporation, the negatively charged LAC molecules in the DES displace PSS anions, promoting rearrangement of positively charged PEDOT chains. This interaction was confirmed in our previous work^55^ through a red shift in the symmetric SO_3_^−^ stretching vibration from 1099 cm^−1^ to 1014 cm^−1^, consistent with PSS displacement, a phenomenon also reported in PEDOT:PSS formulations with cholinium lactate-based ionic liquids^38,58^.

The PEDOT:PSS/DES formulation forms a supramolecular gel that increases viscosity without phase separation^34,59^. When processed *via* spin coating, it enables significantly thicker films (451-818 nm, Figure 1b) compared to conventional PEDOT:PSS/EG/DBSA formulations (hereafter referred to as PEDOT:PSS). To evaluate this advantage, we optimised spin-coating parameters and deposition timing to avoid premature gelation. Using a spin speed of 500 rpm and a high-acceleration ramp (5000 rpm/s), we achieved PEDOT:PSS/DES film thicknesses higher than 800 nm in a single step, substantially exceeding the 117-199 nm range of PEDOT:PSS films under the same conditions. However, the absence of a surfactant and the gel-like nature of the DES formulation resulted in higher thickness variability across the wafer, particularly towards the edges.

Despite the simpler additive composition (13 mg DES vs. 100 mg EG + 2.5 mg DBSA in the standard blend), PEDOT:PSS/DES yielded films 3-5 times thicker, highlighting the importance of supramolecular interactions and secondary doping effects. This enhanced thickness is particularly beneficial in bioelectronic interfaces, where the volumetric capacitance of PEDOT:PSS scales with film thickness, improving electrochemical performance^26,60^. While thicker PEDOT:PSS films can be achieved *via* sequential layering, weak interlayer adhesion may compromise long-term stability^21,26^. In contrast, PEDOT:PSS/DES enables thick, stable films in a single step, making it attractive for applications requiring both conformability and robust electrochemical properties.

To demonstrate applicability, we integrated both PEDOT:PSS/DES and conventional PEDOT:PSS into flexible PaC-based cuff implants for sciatic nerve interfacing in rodents (Figure 1c). The electrode array consisted of 18 circular electrodes (100 μm diameter, 175 μm inter-electrode spacing) and six rectangular electrodes (800 × 120 μm) arranged symmetrically. A 30° pre-formed bend allowed the array to warp around the nerve up to three full turns, accommodating a range of rat nerve diameters (from 200 μm to 1 mm), thereby enabling a generalizable and anatomically adaptive interface design.

The devices were fabricated using standard microfabrication protocols^50^ (Figure 1d) with a Ti/Au metal stack for contact pads and feedlines, and a spin-coated conductive polymer layer for the electrodes. These layers were sandwiched between 2 µm-thick PaC substrate and passivation layers, resulting in a total device thickness of 4 µm. The thin PaC layers ensure that the resulting devices are highly flexible, facilitating efficient wrapping around the nerve. Aiming for maximal conformability and stability, we designed the devices with metal-free electrodes made either from spin-coated PEDOT:PSS/DES or PEDOT:PSS. While the metallic feedlines were patterned using a lift-off technique, the conductive polymer film was spin-coated and patterned *via* dry etching. For PEDOT:PSS/DES, an additional 100 nm thick PaC interlayer was deposited to improve photoresist adhesion. After patterning the conductive polymer layer, the devices were passivated with PaC and structured using a two-step RIE process. The outline of the devices and the contact pad openings were etched first, followed by the etching of the electrode openings.

Final devices (Figure 1e, f) showed a clear visual difference between the thicker PEDOT:PSS/DES (darker blue) and thinner PEDOT:PSS films. Finally, FIB cross-sections confirmed thicknesses of 550 nm for PEDOT:PSS/DES and 171 nm for PEDOT:PSS at the edges of the wafer (Figure 1g, h), validating the enhanced deposition performance of the DES formulation.

### Enhanced electrochemical and electrical performance of PEDOT:PSS/DES electrodes

We carried out an extensive electrochemical and electrical characterisation to evaluate the suitability of the PEDOT:PSS/DES blend for neural recording and stimulation, comprising EIS, estimation of the CSC and CIC, as well as resistivity measurements using the Van der Pauw method. To account for scaling effects in the electrochemical performance, we fabricated test probes featuring different electrode diameters, ranging from 25 µm to 100 µm (Figure S1).

Figure 2a depicts a Bode plot comparing the impedance and phase of PEDOT:PSS/DES and PEDOT:PSS electrodes with a diameter of 100 µm. Overall, PEDOT:PSS/DES demonstrates an excellent impedance with a typical behaviour featuring a resistive regime at high and intermediate frequencies (100 Hz - 100 kHz) where the phase is close to 0° and the impedance is dominated by the access resistance, influenced by the resistive contribution of the electrolyte at the electrode-electrolyte interface. Subsequently, a transition to a capacitive regime at lower frequencies occurs^61^. Interestingly, the PEDOT:PSS electrodes exhibit an additional peak in phase in the frequency range between 100 - 300 Hz. This nontypical behaviour likely originates from oxidation of the top layer of the films during RIE of the contact openings, which was performed with strong oxygen-containing plasma. Due to the larger thickness of the PEDOT:PSS/DES the relative effect of the oxidation is not as prominent and does not result in the formation of a distinct peak. Nevertheless, for both PEDOT:PSS-based materials the phase of the electrodes approaches −90° at low frequencies (below 100 Hz), approximating a perfectly capacitive behaviour as reported before^50,52,62^.

**Figure 2.**
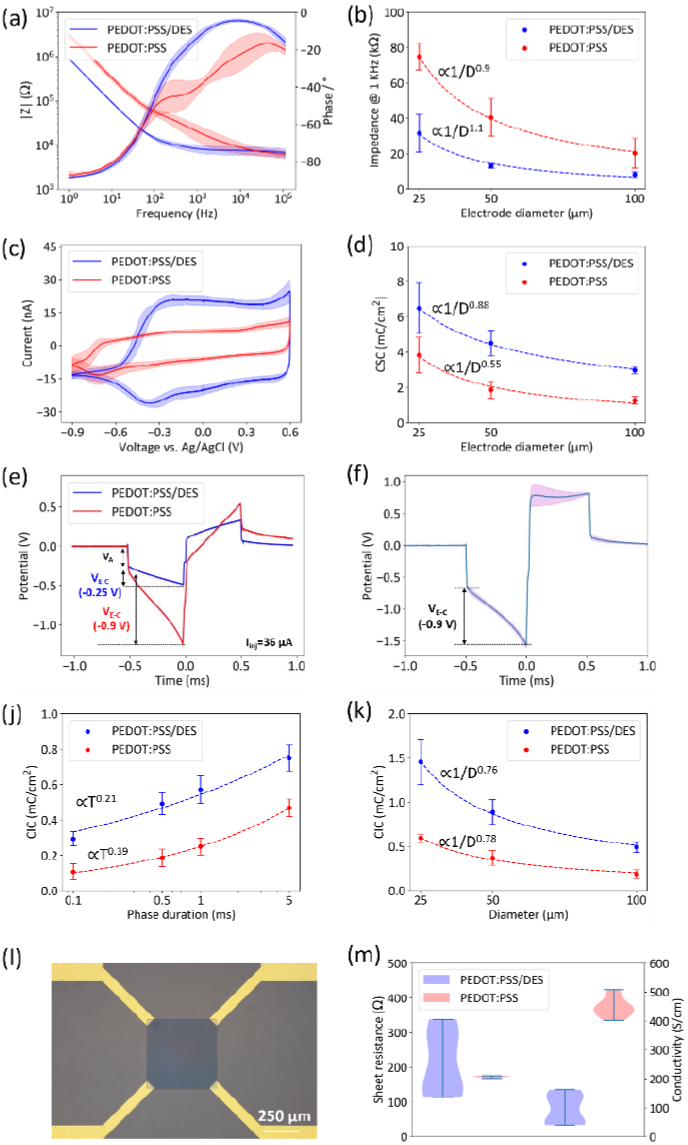
Electrochemical and electrical characterization. (a) Bode plots showing the impedance of 100 µm PEDOT:PSS/DES and PEDOT:PSS electrodes obtained *via* EIS. The shaded region represents the standard deviation (*N*=19). Impedance magnitude at 1 kHz for PEDOT:PSS/DES and PEDOT:PSS electrodes with different diameters (25, 50, and 100 µm). The quality of the fits (*a/D*^*b*^, where *D* is the electrode diameter) is *r*^*2*^=0.985 (blue) and *r*^*2*^=0.999 (red). Cyclic voltammetry of 100 µm PEDOT:PSS/DES (*N*=14) and PEDOT:PSS (*N*=17) electrodes with standard potential limits of −0.9 V and 0.6 V. (d) *CSCc* scaling dependency (*a/D*^*b*^) for PEDOT:PSS/DES (*r*^*2*^=0.983) and PEDOT:PSS (*r*^*2*^=0.999) electrodes of different sizes. (e) Exemplary voltage transient measurements for a PEDOT:PSS and a PEDOT:PSS/DES electrode (100 µm diameter) after applying a squared biphasic current pulse with an amplitude of 36 µA and a pulse width of 0.5 ms. The cathodic voltage excursion of the electrodes (*V*_*E-C*_) is determined after subtracting the access voltage *V*_*A*_. The transient curves were averaged over 10 pulse repetitions (the standard deviation depicted as shaded region). (f) Averaged voltage transient of the 100 µm PEDOT:PSS/DES electrodes (*N*=14) for a pulse width of 0.5 ms at the cathodic excursion limit (*V*_*E-C*_=-0.9 V). The shaded region depicts the standard deviation. (j) CIC pulse width dependency (*a*·*T*^*b*^, where *T* is the pulse width) for PEDOT:PSS/DES (*r*^*2*^=0.971) and PEDOT:PSS (*r*^*2*^=0.999) electrodes (100 µm diameter). (k) *CIC* electrode diameter scaling dependency (*a/D*^*b*^) for PEDOT:PSS/DES (*r*^*2*^=0.998) and PEDOT:PSS (*r*^*2*^=0.992) determined for a fixed pulse width of 0.5 ms. (l) Optical microscopy image of the Van der Pauw structure for conductivity measurements. Metal pads (Ti/Au, yellow) are contacting the edges of the PEDOT:PSS square (dark blue) with a side width of 500 µm. (m) Sheet resistance and conductivity of the PEDOT:PSS/DES and PEDOT:PSS films obtained from multiple measurements (*N*=10).

PEDOT:PSS/DES features a significantly reduced impedance compared to PEDOT:PSS for frequencies below 10 kHz (Figure 2b). The relevant frequency spectrum in the context of neural interfacing depends on the type of recorded signals. While slow signals such as local field potentials (LFPs) are typically analysed at frequencies under 300 Hz, the dominant frequency components of faster action potential signals lie in the range of 300 Hz to 3 kHz^63^. Therefore, neural implants are often benchmarked considering their total impedance value at 1 kHz^61^. The PEDOT:PSS/DES electrodes significantly outperform PEDOT:PSS with respect to this metric, featuring an impedance of 8.3±1.6 kΩ compared to 20.3±8.4 kΩ for PEDOT:PSS, considering a diameter of 100 µm (*p*<0.001, *N*=19). Normalizing the impedance to the GSA yields a specific impedance of 0.652±0.126 Ωcm^2^ and 1.59±0.66 Ωcm^2^ for PEDOT:PSS/DES and PEDOT:PSS, respectively. In addition, PEDOT:PSS/DES electrodes retain their resistive behaviour at lower frequencies, which can facilitate high-fidelity recording of slower signals with better SNR and less signal distortion due to phase shifts. For instance, the reduced impedance would translate to a thermal noise level of 2.02±0.17 µV for PEDOT:PSS/DES and 3.74±0.62 µV for PEDOT:PSS in the low frequency band of 1-300 Hz, and 1.79±0.182 µV and 2.37±0.54 µV in the higher frequency band of 300 Hz – 3 kHz. The enhanced recording capabilities of PEDOT:PSS/DES electrodes are also maintained for smaller electrode dimensions (Figure 2b). For instance, 25 µm PEDOT:PSS/DES electrodes feature an impedance of 31.5±10.6 kΩ, which is still comparable to the impedance of the large PEDOT:PSS electrodes. These results highlight the potential of PEDOT:PSS/DES for enhancing the resolution of neural interfaces, as it can enable the utilisation of smaller electrodes without sacrificing the quality of the recording.

The lower impedance of PEDOT:PSS/DES originates from the high specific capacitance of the electrodes. To estimate the capacitance, we utilized the low frequency spectrum of the EIS data and fitted a simple Randles circuit to model the electrode-electrolyte interface^26^. This resulted in a specific capacitance of 2.22±0.05 mF/cm^2^ for PEDOT:PSS/DES electrodes and 0.65±0.09 mF/cm^2^ for PEDOT:PSS. Taking into account the film thickness as estimated from the FIB cuts, the equivalent volumetric capacitance *C\** amounts to 40 F/cm^3^ and 38 F/cm^3^ for PEDOT:PSS/DES and PEDOT:PSS, respectively. These values are on the lower range compared with existing literature on PEDOT:PSS. For instance, Rivnay *et al*.^64^ reported a *C\** of 39 F cm^−3^ while the value estimated by Biachni et al.^60^ reached 170 F cm^−3^. However, it should be noted that the calculated values likely underestimate the actual volumetric capacitance, since the electrode coating was thinned down during RIE, especially in the centre of the electrodes. This effect is difficult to quantify, as the amount of over-etching depends on the position of the devices on the wafer. Nevertheless, these results suggest that the volumetric capacitance of PEDOT:PSS/DES films is comparable to conventional PEDOT:PSS, and the enhanced electrochemical performance rather stems from its capability of creating thicker electrode films.

To further elucidate the electrochemical characteristics of the materials, we performed CV with potential limits of −0.9 V and 0.6 V, representing the water-window for PEDOT-based electrodes^65^ (Figure 2c). Both PEDOT:PSS/DES and PEDOT:PSS electrodes feature a nearly rectangular shape of the voltammogram, highlighting the pseudocapacitive property of the films. A prominent cathodic peak can be observed for PEDOT:PSS at a potential of roughly −0.7 V, which has been attributed to the injection of electrons in the PEDOT coupled with incorporation of cations into the film^66^. Notably, this peak is shifted to a lower potential of −0.4 V in the case of PEDOT:PSS/DES. The cathodic charge storage capacity (*CSCc*) was calculated for both materials by integrating the cathodic charge in the CV curve, revealing an increase in the *CSCc* by a factor of 2.3 for 100 µm PEDOT:PSS/DES (2.98±0.19 mC/cm^2^) electrodes compared to PEDOT:PSS (1.27±0.21 mC/cm^2^) electrodes. The *CSCc* furthermore increased significantly with the reduction in electrode diameter, reaching 6.48±1.43 mC/cm^2^ for PEDOT:PSS/DES and 3.83±1.01 mC/cm^2^ for PEDOT:PSS, comparable to other works^54^.

Since neural stimulation employs fast pulses in the micro- to millisecond range, only a small fraction of the *CSC* can be safely delivered during a stimulation pulse^67^. We therefore determined the *CIC* of the materials by recording the voltage excursions of the electrodes during biphasic current stimulation using symmetric square charge-balanced pulses with a leading cathodic polarity. Figure 2e depicts the voltage transient of a 100 µm PEDOT:PSS electrode while applying a 0.5 ms current pulse with an amplitude of 36 µA. The maximum cathodic voltage excursion reaches the cathodic limit of −0.9 V, indicating that the electrode is already operating at the current injection limit. In contrast, a PEDOT:PSS/DES electrode with the same size is polarized only up to −0.25 V while injecting the same amount of current. This suggests that PEDOT:PSS/DES electrodes can deliver the same amount of charge at reduced potentials which translates into improved safety and lower energy requirements for neural stimulation.

On average, the PEDOT:PSS/DES electrodes demonstrated a limiting current of 77.3±9.7 µA before reaching the cathodic polarization limit (Figure 2f) which corresponds to a *CIC* of 0.49±0.06 mC/cm^2^, about 2.5 times higher compared to PEDOT:PSS (0.19±0.05 mC/cm^2^). The *CIC* increased consistently for longer phase durations, exceeding 0.7 mC/cm^2^ at 5 ms for 100 µm PEDOT:PSS/DES, compared to just over 0.3 mC/cm^2^ for a 100 µs pulse width. This behaviour is consistent with previous findings^54^ and suggests that prolonged phase durations permit higher charge densities. Similarly to the *CSC*, the CIC also exhibits a strong dependency on the electrode size, reaching 1.46±0.26 mC/cm^2^ and 0.6±0.04 mC/cm^2^ at 25 µm for PEDOT:PSS/DES and PEDOT:PSS, respectively. It should be noted that the *CIC* values for PEDOT:PSS are relatively low compared to other reports^54^, however these discrepancies can be attributed to a combination of factors, such as variations in the film thickness, differences in the processing of the film (*i*.*e*., dry etching vs. stencil patterning), over-etching of the film, as well as to the absence of metal contacts under the electrodes. For instance, Cho *et al*.^68^ recently reported a *CIC* of 0.7 mC/cm^2^ for 0.4 ms pulses using bare PEDOT:PSS electrodes, which closely resembles the values found in this work. The size-dependent *CIC*, on the other hand, can be related to edge effects^54^, which are likely exacerbated by the absence of a highly conductive metallic surface under the electrodes, as well as to the contribution of the PEDOT:PSS feedline, which was kept at a constant width for all electrode sizes.

Finally, the conductivity of the films was estimated using the Van der Pauw method. To this end, we utilized dedicated structures featuring gold pads contacting the edges of a 500 µm square of conductive film (Figure 2l). The measurements yielded a conductivity of 447.3±32.8 S/cm for the standard PEDOT:PSS blend, in good agreement with other reports on PEDOT:PSS films with a similar composition^50^. In this regard, the PEDOT:PSS outperformed the novel PEDOT:PSS/DES blend, which proved to be roughly 5 times less conductive (94.6±51.3 S/cm). The latter is expected, given that additives such as EG and DBSA have been shown to increase the conductivity of PEDOT:PSS films, whereas GOPS has the opposite effect^69^, resulting in films that are less conductive than PEDOT:PSS/DES blends without GOPS, as reported privously^55^. Additionally, we observed a large variability in the film conductivity across the wafer, suggesting that some inhomogeneities in the deposited film might be present. Nevertheless, the sheet resistance of the PEDOT:PSS/DES was comparable to PEDOT:PSS (211.3±91.9 Ω/sq vs. 172.8±2.44 Ω/sq), indicating that the increased thickness of the PEDOT:PSS/DES film is sufficient to compensate for the reduced conductivity.

### Enhanced electrochemical properties of PEDOT:PSS/DES translate to superior signal recording and stimulation efficiency

To evaluate the functional relevance of the enhanced electrochemical properties observed *in vitro*, we performed *in vivo* experiments using the cuff multielectrode arrays described in the previous section, comparing PEDOT:PSS/DES and conventional PEDOT:PSS electrodes for neural recording and stimulation of the rat sciatic nerve in non-recovery implantations (Figure 3a). PEDOT:PSS/DES electrodes demonstrated superior performance compared to PEDOT:PSS across multiple electrophysiological metrics, including impedance characteristics, signal acquisition, and stimulation efficiency.

**Figure 3.**
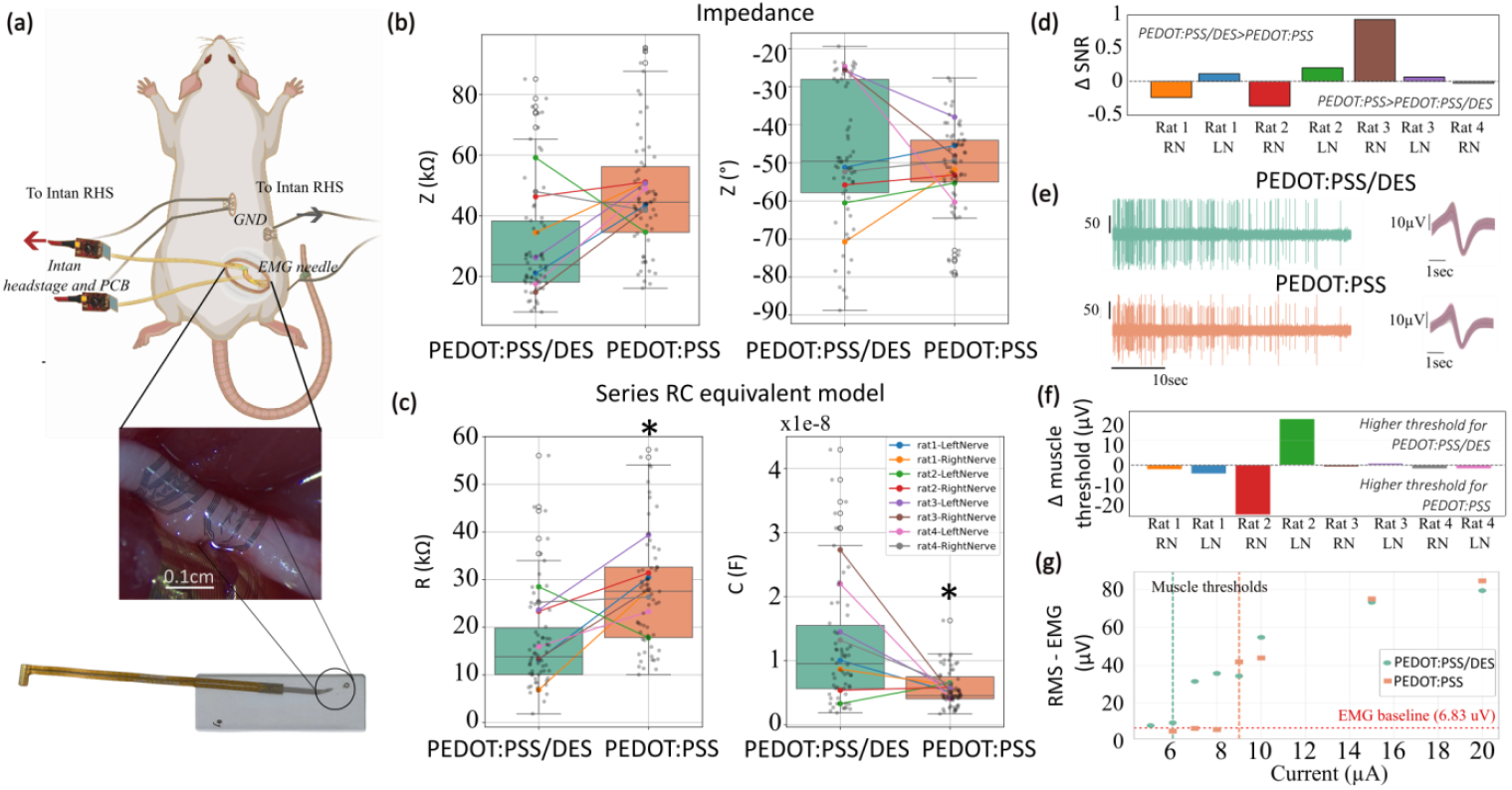
In vivo evaluation of PEDOT:PSS and PEDOT:PSS/DES devices. (a) Schematic of the experimental setup showing surgical implantation at the sciatic nerve with side-by-side placement of the two cuffs. Inset: magnified view of the cuff placement on the sciatic nerve. Bottom: photograph of the bonded electrode assembly with FFC connector interfacing with a custom-made PCB, which connects to the Intan RHS headstage. EMG needle electrodes were placed in the tibialis anterior muscle. All signals were recorded via the Intan RHS system. (b) Impedance magnitude and phase at 1 kHz for PEDOT:PSS/DES (green) and PEDOT:PSS (blue). (c) Resistance and capacitance values extracted from equivalent series RC circuit modeling, revealing statistically significant differences between materials (paired t-test, p < 0.05). (d) Trial-wise difference in signal-to-noise ratio (SNR) between materials (PEDOT:PSS/DES – PEDOT:PSS) for both right and left sciatic nerves across four rats (8 independent trials). (e) Representative neural recordings from PEDOT:PSS/DES and PEDOT:PSS during evoked pain activity, with corresponding average spike waveforms. (f) Differences in stimulation threshold (minimum current to evoke a visible motor twitch) between materials. (g) Representative EMG RMS responses at increasing stimulation currents for each device. Each point represents the average RMS over 20 stimulation pulses (pulse duration: 100 µs). Red dashed line represents EMG RMS at baseline prior to stimulation. Vertical dashed lines represents the currents at which a motor response was visually observed.

Impedance magnitude and phase values extracted from baseline recordings did not differ significantly between materials (Figure 3b). However, PEDOT:PSS/DES consistently exhibited lower impedance magnitude across most animals. When modelled using a series RC circuit, implanted PEDOT:PSS/DES electrodes showed significantly lower resistance and higher capacitance than PEDOT:PSS electrodes (Figure 3c), indicating improved charge transfer and capacitive behaviour. The influence of stray capacitance from the experimental setup cannot be excluded and may affect impedance measurements. As a result, direct comparisons between *in vitro* and *in vivo* values are not encouraged. Here, all *in vivo* comparisons were performed between paired cuffs implanted simultaneously in the same animal to ensure consistency and control for inter-animal variability.

The improved electrochemical and conformal properties of PEDOT:PSS/DES translated into enhanced neural signal acquisition (Figure 3d). SNR was higher for PEDOT:PSS/DES in the majority of trials, suggesting greater neural recording quality (Figure 3d and e), although this trend did not reach statistical significance using a paired *t*-test. Only two trials showed superior performance for PEDOT:PSS, one of which exhibited similar impedance and capacitance across materials. This suggests that while the materials’ electrochemical and mechanical properties are important, they are not the sole determinant of the recording and stimulation efficacy. Other factors, such as the proximity of the electrodes to motor and sensory fascicles, play a significant role in shaping functional outcomes. Additionally, spike amplitude was generally greater, although not significantly in statistical terms, with PEDOT:PSS/DES in five out of seven trials, further supporting its enhanced recording capability (Figure 3e).

Stimulation responses, as assessed by EMG RMS amplitude across increasing current levels, were broadly similar between PEDOT:PSS/DES and PEDOT:PSS electrodes. However, PEDOT:PSS/DES more reliably elicited visible motor responses at lower current thresholds, indicating improved recruitment of the tibialis anterior muscle (Figure 3f). This advantage was generally not reflected in EMG signal magnitude at matched currents (Figure in S6), where statistical comparisons at 10 µA and 30 µA stimulation currents using paired *t*-tests did not reveal significant differences between materials (*p* = 0.05). However, clearer distinctions emerged in some cases at lower current levels (Figure 3g, corresponding to Rat 1-RN).

The improved stimulation efficacy of PEDOT:PSS/DES is likely related to its enhanced electrochemical properties, including lower series resistance and higher capacitance, which support more efficient charge delivery and better coupling at the electrode-tissue interface. These characteristics may reduce the current required to reach activation thresholds, even when EMG output remains comparable.

Notably, the only case in which PEDOT:PSS outperformed PEDOT:PSS/DES corresponded to a situation where PEDOT:PSS had clearly superior electrochemical properties (higher capacitance and lower impedance), consistent with expected trends. However, in the case where PEDOT:PSS/DES showed the largest functional advantage, the difference in electrochemical performance compared to PEDOT:PSS was relatively modest. In contrast, other recordings with more dramatic electrochemical differences showed smaller functional effects, indicating, once again, that local interface conditions may influence performance. Comparing the magnitude of differences across trials is not advised, as each trial may involve slightly different electrodes in terms of number and placement. It is more appropriate to focus on differences between materials within a single trial, where electrode configuration and placement remain more consistent.

## Conclusions

This study demonstrates that incorporating DES (ChCl:LAC) into PEDOT:PSS formulations offers a powerful strategy to enhance the electrochemical performance of implantable bioelectronic interfaces. We developed a PEDOT:PSS/DES formulation that facilitates the processing of thick, conductive polymer, gel-like film electrodes, compatible with established microfabrication workflows based on thin-film technology. The enhanced thickness of PEDOT:PSS/DES electrodes resulted in exceptional electrochemical characteristics, paving the way for miniaturized, higher-density neural interfaces without sacrificing functionality.

The use of eutectogels opens a new venue in the tailored modification of PEDOT:PSS, such as anticorrosion resistance, mixed ionic-electronic conductivity, and long-term mechanical stability, features that are critical for chronic implantation and multi-modal bioelectronic therapies. Future work will focus on evaluating these properties under long-term electrophysiology setups, and the use of alternative DES chemistries to further expand the functional landscape of PEDOT-based materials for implantable technologies.

## Author contributions

V.R.M and A.G.G supervised and contributed equally to this work. M.K, R.R.M, A.D.A, V.R.M, and A.G.G conceived and planned the study. R.R.M and A.D.A developed the PEDOT:PSS/DES formulation. A.D.A, M.L.P, and D.M provided the DES. M.K and V.R.M established the microfabrication process-flow, fabricated the probes, and carried out the electrochemical and electrical characterisations. M.K processed the data of *in vitro* characterisations with the support of V.R.M. A.G.G and C.L carried out the *in vivo* validation. A.G.G developed the custom code and processed the *in vivo* data. D.M. and G.G.M. contributed laboratory facilities, equipment, and other resources required for the experimental work. M.K, R.R.M, A.D.A, V.R.M, and A.G.G wrote the initial draft of the manuscript. All authors reviewed and edited the manuscript.

## Data availability

The data supporting the electrochemical characterisation have been included as part of the Supplementary Information. All data collected for the animal work, including the recording and stimulation electrophysiology datasets, are available at Zenodo public repository with DOI: https://doi.org/10.5281/zenodo.16746532 (under embargo until accepted for publication). The code for analysing the electrophysiology data can be found at GitHub with DOI: 10.5281/zenodo.16763365. The version of the code employed for this study is version V1.

## Acknowledgements

The authors thank the Helmholtz Nano Facility (HNF) at *Forschungszentrum* Jülich for facilitating the microfabrication and S. Decke, E. Yilmaz, and R. Stockmann for providing support during microfabrication. The authors also thank E. Brauweiler-Reuters for carrying out FIB sectioning and SEM. The authors thank A. Offenhäusser for infrastructural and scientific support. R.R.M acknowledges support from EPSRC grant [EP/S022139/1]. G.G.M. and A.D.A acknowledge support from EU grant COPE-Nano (ID: 101059828). M.L.P and D.M gratefully acknowledge the financial support from IKERBASQUE-Basque Foundation for Science and the Marie Sklodowska-Curie Research and Innovation Staff Exchanges program under grant agreement IONBIKE 2.0 MSCA-SE 101129945. A.G. acknowledges support from the Rosetrees Trust and the Royal Academy of Engineering. This work was funded by UKRI grants and was supported by the Deutsche Forschungsgemeinschaft (DFG, German Research Foundation; GRK2610 (project number 424556709). For the purpose of open access, the author has applied a Creative Commons Attribution (CC BY) licence to any Author Accepted Manuscript version arising.

