## Supplementary information for "PEDOT:PSS conducting eutectogel for enhanced electrical recording and stimulation in implantable neural interfaces"

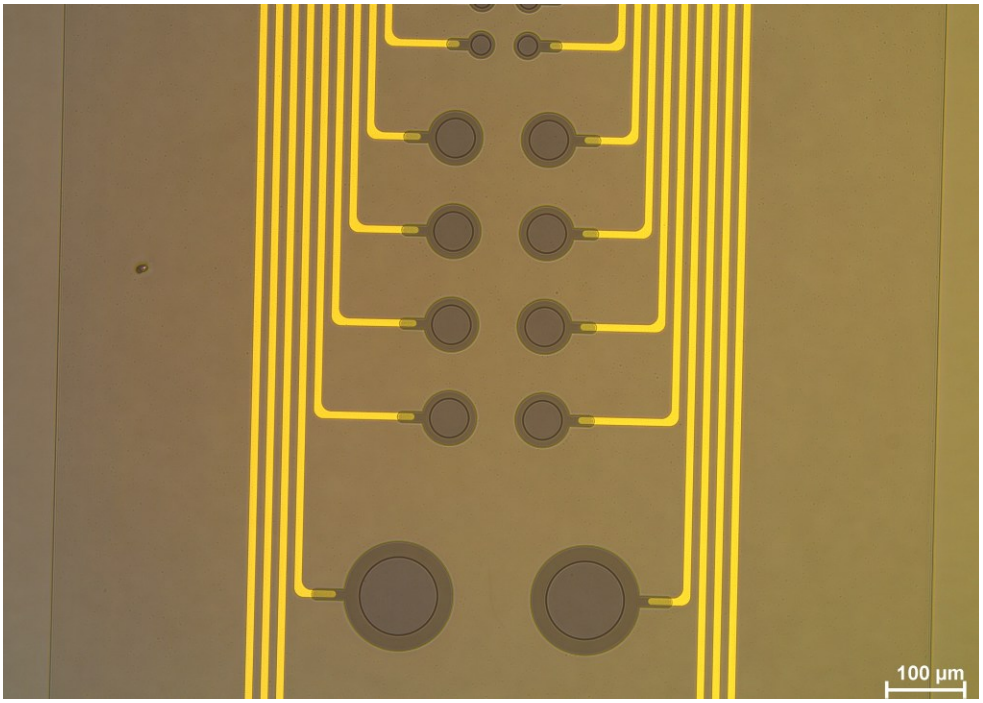


**Figure S1. Design of characterization probes.** Three electrode diameters were used for testing the scaling effects on the electrochemical performance of PEDOT:PSS/DES and PEDOT:PSS probes. The picture showcases a PEDT:PSS/DES probe with electrode diameters of 25 µm (top rows), 50 µm (middle rows), and 100 µm (lower row).

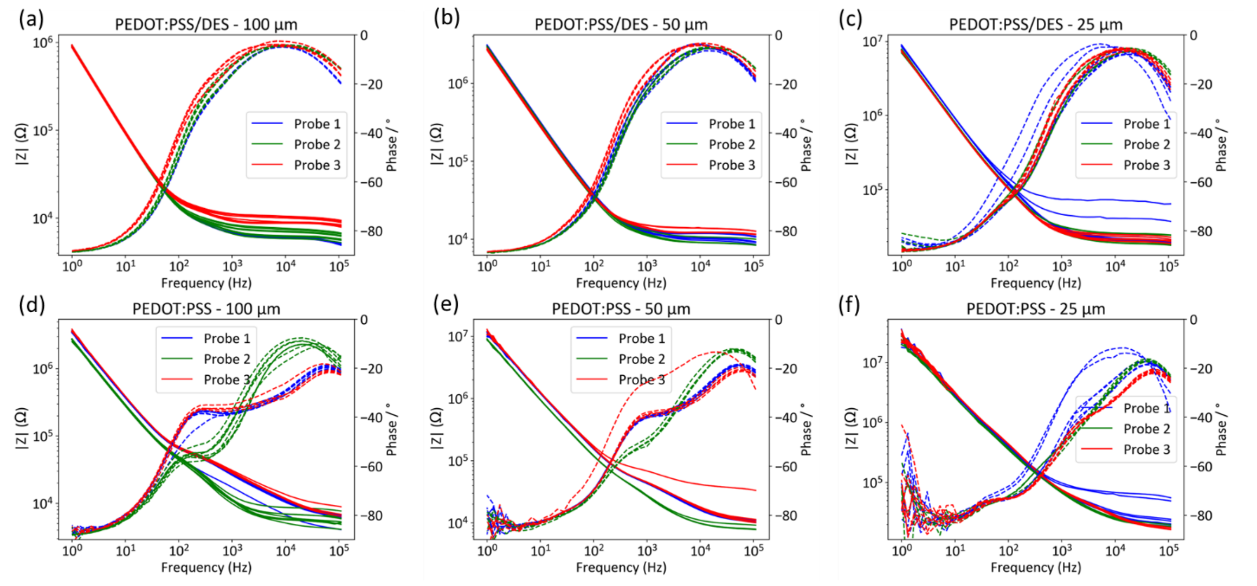


**Figure S2. Raw data of electrochemical impedance spectroscopy measurements**. Bode plots depicting the impedance magnitude ($|Z|$, left axis) and phase (right axis). Upper row (a-c): PEDOT:PSS/DES electrodes with a diameter of 100 µm (*N*=19), 50 µm (*N*=18) and 25 µm (*N*=21). Lower row (d-f): PEDOT:PSS electrodes with a diameter of 100 µm (*N*=19), 50 µm (*N*=17) and 25 µm (*N*=18).

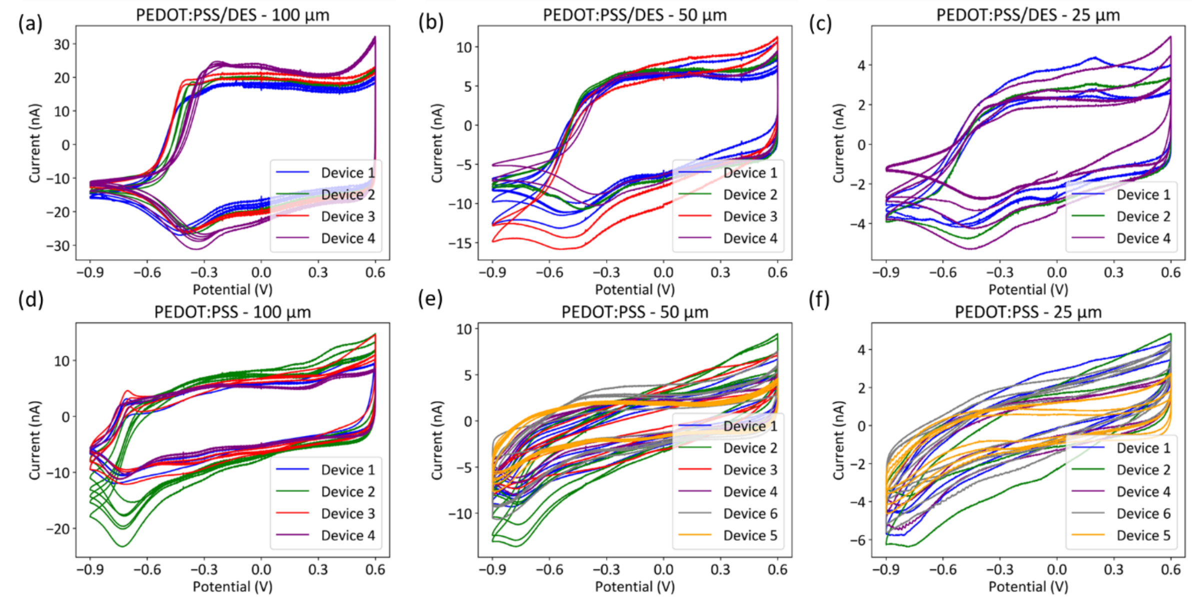


**Figure S3. Raw data of cyclic voltammetry measurements.** Cyclic voltammograms measured with a scan rate of 100 mV/s versus an Ag/AgCl reference electrode. Upper row (a-c): PEDOT:PSS/DES electrodes with a diameter of 100 µm (*N*=14), 50 µm (*N*=11) and 25 µm (*N*=8). Lower row (d-f): PEDOT:PSS electrodes with a diameter of 100 µm (*N*=17), 50 µm (*N*=29) and 25 µm (*N*=16).

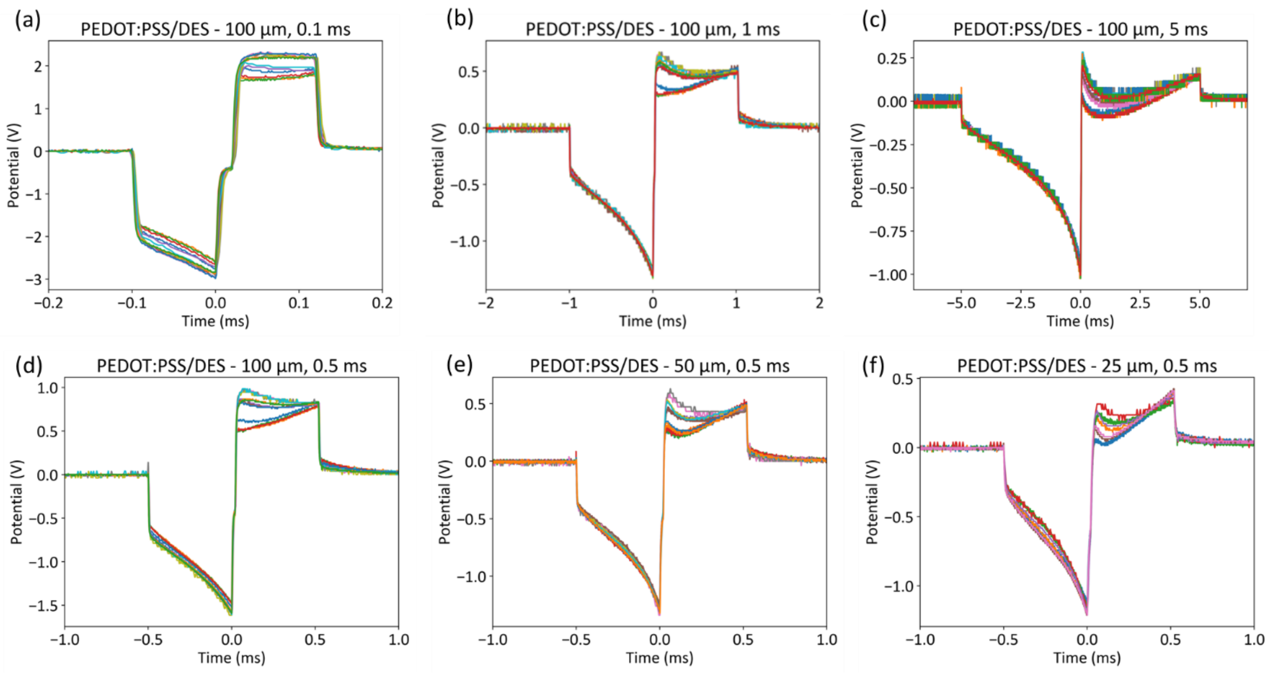


**Figure S4. Voltage transients for PEDOT:PSS/DES electrodes recorded at the current injection limit**. The maximum cathodic voltage excursion ($V_{E\text{-}C}$) reaches the cathodic limit of -0.9 V. For all electrodes and conditions, the cathodic limit was reached before the anodic limit. Upper row (a-c): Voltage transients of electrodes with a diameter of 100 µm and a pulse width of 0.1 ms (*N*=13), 1 ms (*N*=13), and 5 ms (*N*=13). Lower row (d-f): Voltage transients for a pulse width of 0.5 ms and electrode diameters of 100 µm (*N*=13), 50 µm (*N*=12) and 25 µm (*N*=7).

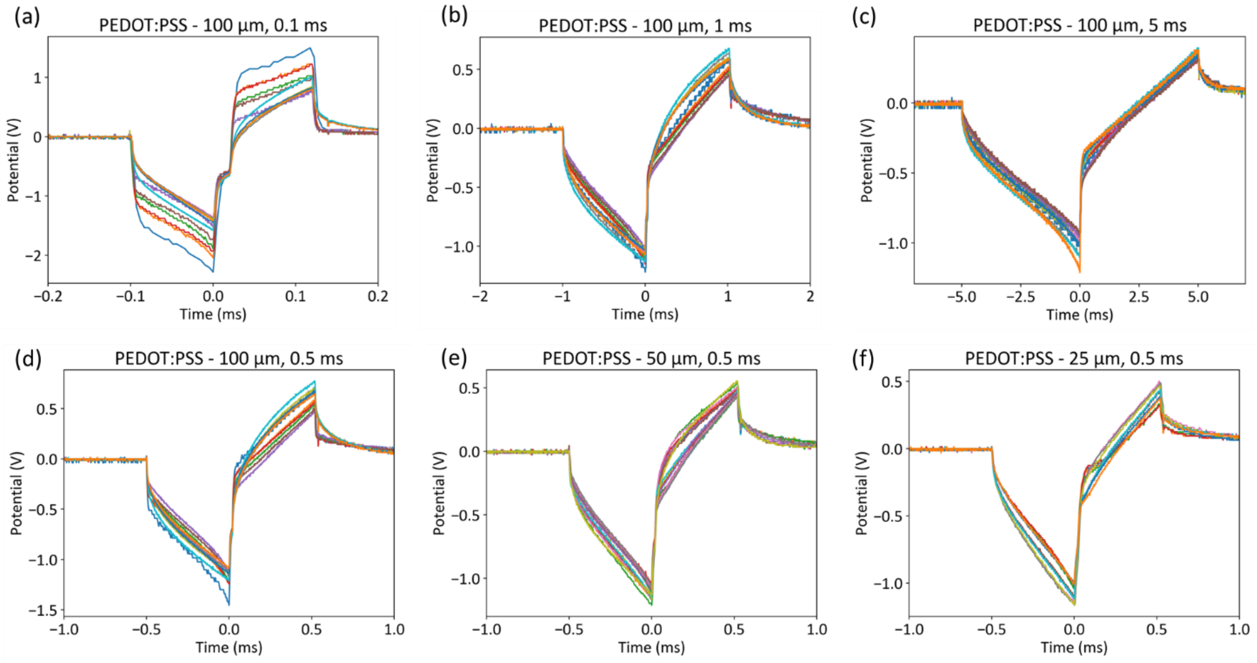


**Figure S5. Voltage transients for PEDOT:PSS electrodes recorded at the current injection limit.** The maximum cathodic voltage excursion ($V_{E\text{-}C}$) reaches the cathodic limit of -0.9 V. For all electrodes and conditions, the cathodic limit was reached before the anodic limit. Upper row (a-c): Voltage transients of electrodes with a diameter of 100 µm and a pulse width of 0.1 ms (*N*=12), 1 ms (*N*=12) and 5 ms (*N*=12). Lower row (d-f): Voltage transients for a pulse width of 0.5 ms and electrode diameters of 100 µm (*N*=12), 50 µm (*N*=19) and 25 µm (*N*=12).

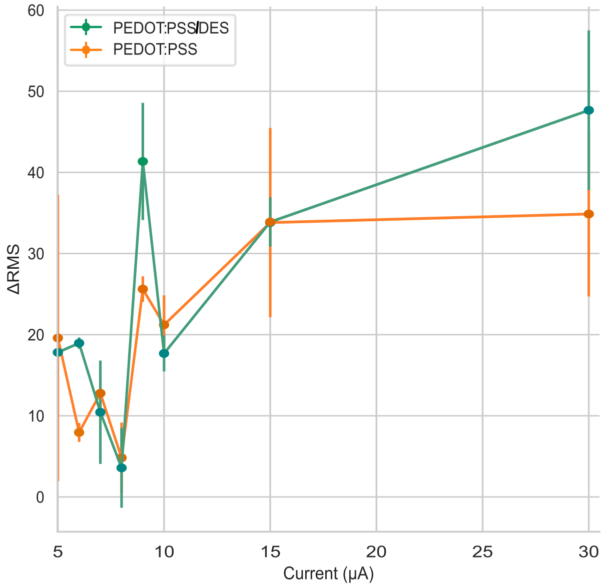


**Figure S6. Relative increase in EMG RMS amplitude compared to baseline at each stimulation current.** Pulse width at 100 µs. Each data point represents the average response from 20 stimulation pulses, included only when at least four trials were available for that stimulation condition. No statistical significance was found in differences at 10 µA or 30 µA (paired *t*-test, $p = 0.05$).
